# GrAnnoT, a tool for efficient and reliable annotation transfer through pangenome graph

**DOI:** 10.1101/2025.02.26.640337

**Authors:** Nina Marthe, Matthias Zytnicki, Francois Sabot

**Affiliations:** DIADE unit, UM, Cirad, IRD, 911 avenue Agropolis Montpellier F-34391, France; MIAT, INRAE, Auzeville-Tolosane F31320, France

**Keywords:** Pangenome, graph, annotation transfer

## Abstract

The increasing availability of genome sequences has highlighted the limitations of using a single reference genome to represent the diversity within a species. Pangenomes, encompassing the genomic information from multiple genomes, thus offer a more comprehensive representation of intraspecific diversity. However, pangenomes in form of a variation graph often lack annotation information and tools for, which limits their utility for downstream analyses. We introduce here GrAnnoT, a tool designed for an efficient and reliable integration of annotation information in such variation graphs. It projects existing annotations from a source genome to the variation graph and subsequently to other embedded genomes. GrAnnoT was benchmarked against state-of-the-art tools on pangenome variation graphs and linear genomes from Asian rice, and tested on human and *E. coli* data. The results demonstrate that GrAnnoT is consensual, conservative, and fast. It provides informative outputs, such as presence-absence matrices for genes, and alignments of transferred features between source and target genomes, helping in the study of genomic variations and evolution. GrAnnoT’s robustness and replicability across different species make it a valuable tool for enhancing pangenome analyses. GrAnnoT is available under the GNU GPLv3 licence at https://forge.ird.fr/diade/dynadiv/grannot.

## Introduction

Recent advances in genome sequencing and assembly methods give access to a massive and increasing number of genome sequences per species for the scientific community. Consequently, while still currently prevalent, the use of a single reference genome has been shown to bias many analyses (Chen et al., 2021; Martiniano et al., 2020; Maurstad et al., 2024), as it favors variant calling toward reference alleles and thus hinders the identification of non-reference sequences. From that rises the idea that a single individual is not enough to represent the diversity of a given species or group. This led to the development and diffusion of the concept of pangenomics across the whole tree of life (Bayer et al., 2020; Liao et al., 2023; Miga and Wang, 2021; Rouli et al., 2015; Shi et al., 2023).

A pangenome aims to represent the complete genomic information from several genomes of the same species or group, in order to better represent the intra-specific/group diversity. While the concept emerged from bacterial studies (Tettelin et al., 2005), it is now applied to larger and more complex eukaryotic genomes. Many studies in pangenomics have been published, and allowed a better understanding of genomic diversity, population dynamics, and evolution (Rice et al., 2023; Secomandi et al., 2025; Tranchant-Dubreuil et al., 2019; Zhou et al., 2022).

This pangenomic information can be stored in different structures depending on the type of organism and study involved. Ranging from gene set to whole pangenome graphs, the methods to build, manipulate and study these structures differ. Some representations of pangenomes have an extensive toolset allowing in depth analysis (e.g. bacterial pangene set with tools such as PPanGGOLiN (Gautreau et al., 2021)), some can handle thousands of genomes (e.g. de Bruijn graphs built by Bifrost (Holley and Melsted, 2020)), and some others still suffer from methodological shortage.

Annotation is an important element for studying genomic sequences and understanding their potential biological functions, if any, to help interpret the variations found in the pangenome in regard of phenotypes. Numerous tools have already been proposed to produce, cluster, visualize, or manipulate pangenome annotation (Durant et al., 2021; Gautreau et al., 2021; Horsfield et al., 2023; Pedersen et al., 2016).

The variation graph (Outten and Warren, 2021) is a promising structure for representing a pangenome, but it still lacks adapted tools to integrate and manipulate annotation. This structure represents the whole sequence information of the embedded genomes, including intergenic regions, small and large variants (SNPs as well as large indels, duplications and translocations - to some extent), and is easier to use for large or complex genomes with high repeat content (Andreace et al., 2023; Secomandi et al., 2025) compared to de Brujin graphs, for instance. In variation graphs, the nodes represent sequences by stretch of DNA, the links (or edges) show the adjacencies between two sequences (nodes) in at least one embedded genome, and the paths reconstruct these embedded genome sequences. Variation graphs are usually built from complete alignments between whole genomes, and the currently most popular tools include PGGB (Garrison et al., 2024) and minigraph-cactus (Hickey et al., 2023). Additionally, tools like VG (Garrison et al., 2018) and ODGI (Guarracino et al., 2022) manipulate these graphs and perform various tasks. These variation graphs are currently used for better alignment of reads, genotyping, and structural variation detection (Hickey et al., 2020; Sirén et al., 2021). However, the variation graph only encodes sequence information, and does not carry any annotation natively. Such biological information is crucial to give context to any variation in genomic sequences, and is often available for the linear reference genomes. Integrating these existing annotations to the variation graphs would enrich these structures and make them a better tool for studying pangenomes.

While tools exist to visualize annotations on a graph (Jonkheer et al., 2022; Liu et al., 2024; Miao and Yue, 2025; Wick et al., 2015), their use is not adapted for large-scale analyses of thousands of annotations. To answer this issue, VG annotate recently proposed to project genomic annotation on a graph, and offers options to efficiently index and query the resulting graph annotation (Novak et al., 2024). However, it cannot project annotations from the graph to the genomes, which would in turn allow to output genome annotation and to identify the variants between the embedded genomes in the annotated regions, to study their impact. Tools for transferring annotation exist for linear to linear genomes, with the most used of them being Liftoff (Shumate and Salzberg, 2021), but it relies on gene-by-gene sequence alignment and does not provide explicit information about variations between genomes. In this regard, the use of the variation graph, which represents in its essence a whole genome alignment and models the synteny between embedded genomes, is a natural way to transfer annotations from genome to graph, or from graph to genome, and to identify the differences between genomes.

To fill this gap, we developed GrAnnoT, a command line tool that manipulates genomic annotations in a variation graph space under its native GFA format. Starting from projecting on the graph the annotation of a single genome in a GFF format, GrAnnoT then outputs the graph annotation in GAF format (as VG annotate), but it also projects this annotation on the other embedded genomes with a dedicated GFF output for each of them. In addition to this transfer, GrAnnoT compares the annotated regions between the genomes in the graph, and outputs transfer statistics (i.e. transfer rate, or mean sequence identity and coverage), lists and types of variants, alignments, and a presence-absence matrix for gene features. These operations rely on the structure of the graph, speeding up the transfers by harnessing the multiple genome alignment it represents (Hickey et al., 2023; Li et al., 2020). We applied it to variation graphs built from different species, and compared it to existing methods to ensure the annotation transfer is valid. As an graph annotation transfer tool, GrAnnoT is fast and conservative, and it allows to study the annotated regions of a pangenome variation graph and of its embedded genomes in an easier way and at a larger scale than any tool before.

## 1. Implementation

GrAnnoT is implemented in Python 3.10+, as a Linux command-line tool that can be installed as a standard python package. It only requires the *tqdm* package and the external program bedtools (Quinlan, 2014) (that must be accessible in the user or in the global path). To ease its installation and use, an AppTainer container definition is available in the GrAnnoT repository on the IRD forge (https://forge.ird.fr/diade/dynadiv/grannot); all the codes are under the GNU GPLv3 license.

### 1.1. Code overview

GrAnnoT performs annotation transfer from an annotated genome (the source genome) to a variation graph (Figure 1). It can also transfer the annotation from the graph to one, several or all other genomes embedded in the graph (the target genomes). It takes as input the annotation of the source genome in GFF3 format, and the variation graph that includes the source and target genome in GFA 1.1 format (without overlap between the nodes).

**Figure 1.**
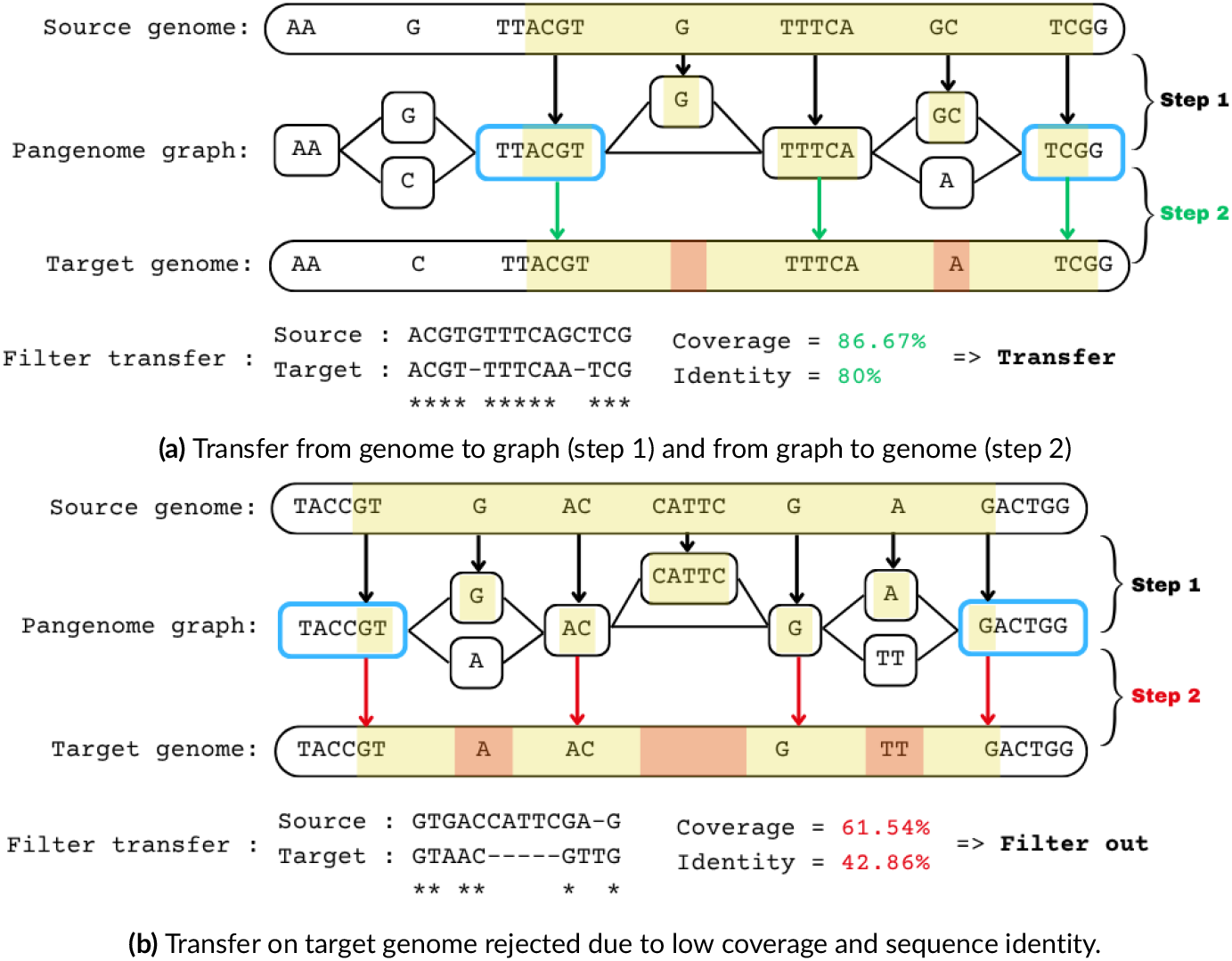
GrAnnoT overview. Step 1: the position of the feature is projected from the source genome to the graph using the positions of the nodes on the source genome. Step 2: the position of the feature is projected from the graph to the target genome (a). The first and last nodes from the feature that are on the target genome (the blue ones) are the ends of the feature in this genome, and everything in between is considered as part of the feature. The differences between the two genomes in this region in terms of path in the graph mirror the differences between the two versions of the feature. If these differences are too important and the transfer does not reach the coverage and/or sequence identity thresholds, the annotation is not transferred (b).

The annotation transfer only relies on the graph structure, harnessing the multiple alignment and synteny it naturally represents. GrAnnoT projects the coordinates between the graph and the genomes, transferring annotations in a fast, alignment-free manner.

Once the annotation has been loaded, GrAnnoT outputs the graph annotation in GAF format (Li et al., 2020). This tab-delimited text format was originally proposed to represent sequence-to-graph alignment. However, it can also be used for graph annotation (Novak et al., 2024), where, instead of describing the paths of the mapped reads, it describes the paths of the annotated features through the graph. GrAnnoT can then output the annotation in GFF3 format for a chosen set of target linear genomes included in the variation graph. These transfers can be filtered through sequence identity and coverage scores similarly to the BLAST approach (Altschul et al., 1990). For these transfers, the alignment of each feature between the source genome and the target one can be outputted in a Clustal-like format, as well as a list of all the variants recorded in the alignments. These alignments are not computed by GrAnnoT, but directly extracted from the graph structure. Finally, a presence-absence matrix for gene features summarizes the transfer on the target genomes.

### 1.2. Implementation details

The first step is to find the start and stop positions of each node from the graph on the embedded genomes (Figure 1, step 1). For that, GrAnnoT follows the paths of these genomes in the graph and computes the start and stop positions of the nodes for each of them; these positions are then stored in BED files, one per contig/chromosome per genome. Then, the BED files representing the source genome are compared to its annotation file using bedtools intersect (Quinlan, 2014). The resulting BED file is processed to compute the paths of the features in the graph and output the graph annotation in the GAF format.

In order to transfer an annotation to a target genome, the sub-path of the genome corresponding to the feature is extracted (Figure 1a, step 2). For that, all the nodes from the original feature path are looked for in the target genome paths. These nodes are then grouped into copies of the feature, since the nodes corresponding to a feature can sometimes be found multiple times in the target genome path (duplication). For each copy, the first and the last nodes are considered as the ends of the feature’s copy in the target genome, and all copies are transferred by default. An option allows to only transfer the copy with the highest sequence identity and coverage. All the nodes between the first and last segment in the target genome path are expected to be part of the feature’s copy to transfer, including the nodes absent from the original feature path, corresponding to insertions. Nodes from the original feature path that are not found in the target genome correspond to deletions. An insertion and a deletion at the same locus in the variation graph correspond to a substitution.

For the transfer itself, only the two nodes at the ends of the feature path on the target genome are considered (nodes in blue in Figure 1). The BED file previously computed reporting the positions of the nodes on the target genome is used to locate these two nodes on the genome.

Transferred features are then filtered based on the coverage (in base) and the identity level between the source and the target genomes (Figure 1b), both set at 80% by default (but can be defined by the user). These parameters are estimated by computing the cumulated length of the shared and different nodes between the paths of the features in the two genomes. The genes are excluded if they do not meet the threshold set, and their child features (exons, CDS, UTR) are not transferred. This filtering ensures that the sequence of the annotated feature is conserved between the source and target genome, to remove spurious transfers. The output is finally printed out in the GFF3 format.

If the user is interested in the differences between the source and target annotation, GrAnnoT can provide a detailed comparison between the feature alternative paths in the source and any of the embedded target genomes. For that, GrAnnoT can output the variants details in a human-readable text format that describes all the variants present in the feature (node deletion, insertion, substitution). A Clustal-like alignment file of all the transferred features based on their alternative paths is similarly generated.

## 2. Benchmark

### 2.1. Data and tools for benchmarks

The main test data used in this paper is an Asian rice pangenome graph built with 13 genomes (Kawahara et al., 2013a; Zhou et al., 2020) using minigraph-cactus v2.8.2 with default options (Hickey et al., 2023; see supplementary data for the exact commands) and the cv Nipponbare IRGSP1.0 as reference. The rice genome is 380-410Mb long and has 12 chromosomes. The annotation used as source (Kawahara et al., 2013b) includes 57,585 *gene* features for 813,790 total features, and is rich in transposable elements (15,848/57,858≈27% of *gene* features are annotated as transposable elements).

GrAnnoT was also tested on a graph of the human chromosome 1 with 92 haplotypes (from Liao et al., 2023) and an *E. coli* 13 genomes graph (see supplementary data for the genomes used) built using the same protocol as for rice (detailed commands available online, Marthe and Sabot, 2025b).

GrAnnoT was compared to existing and state-of-the-art tools (see below) that can also perform annotation transfer in order to assess its efficiency, and using the different data presented before to test its replicability and robustness. All analyses were ran on a biprocessor Intel(R) Xeon(R) CPU E5-2650 v4 @ 2.20GHz with 48 HT CPU computer with 144Gb of RAM, under RockyLinux 9.1 Blue Onyx.

In all transfers made with GrAnnoT, the parameters for sequence identity and coverage based filtering were left as default, so at 80% both. These parameters can be changed by the user, and an evaluation of their effect on the transfers in rice data is available in supplementary data (Table S14).

Multiple tools are available to perform annotation transfer between linear genomes, with different approaches. Tools like CAT (Fiddes et al., 2018), RATT (Otto et al., 2011), FLO (Pracana et al., 2017), or CrossMap (Zhao et al., 2013) use a form of whole genome alignment to convert the positions of the annotated features from one genome to another. Tools like Liftoff (Shumate and Salzberg, 2021), GeMoMa (Keilwagen et al., 2019) and LiftOn (Chao et al., 2024) align the sequence of each annotated feature on the target genome to find its position. The current state-of-the-art annotation transfer tool for linear genome sequences is Liftoff (Shumate and Salzberg, 2021). It is widely used (Alonge et al., 2022; Kim et al., 2021; Wang et al., 2021; Yang et al., 2023), and was chosen here test the validity of GrAnnoT’s genome to genome annotation transfers. However, since Liftoff does not use a pangenome graph to transfer annotations, the comparison with GrAnnoT is biased by the graph itself, whose structure partially impacts the results of GrAnnoT transfer (see Discussion).

For annotation transfer on graph through alignment, Liftoff approach can be mimicked by aligning the sequences of the annotated features to the graph. Graph pangenome alignment tools can be thus compared to GrAnnoT for graph annotation transfer: GraphAligner was chosen for this purpose (Rautiainen and Marschall, 2020), as a state-of-the-art tool for aligning long sequences on a graph.

To transfer annotations between genomes through a pangenome graph and use graph properties instead of gene sequence alignment, VG and ODGI were used. They are state-of-the-art tools for pangenome graph manipulation (Garrison et al., 2018; Guarracino et al., 2022), and while they do not have options specifically designed to transfer annotations between genomes of the graph, they do have options to project coordinates between the graph and its embedded genomes. *odgi position* and *vg inject/surject* project coordinates between the genomes of the graph and can be used to transfer annotations, with the limitation that there is no filtering based on sequence identity of coverage.

Additionally, VG recently implemented an option to annotate the graph by projecting an annotation from a genome to the graph. Although *vg inject* already performed this task, *vg annotate* is specifically designed for this purpose (Novak et al., 2024) and is easier to use. It also offers efficient ways to index an query the graph annotation.

The results and execution time of all these functions were compared to GrAnnoT.

The versions of the tools used are available in the supplementary data. The complete exact commands used for those benchmark are available online (Marthe and Sabot, 2025b). The Jupyter notebooks used for the analysis are available on our Forge (https://forge.ird.fr/diade/dynadiv/grannot, Marthe et al., 2025). All the data used for the analysis and the outputs are available online (Marthe and Sabot, 2025a,b).

### 2.2. Comparison of the transfers

#### Comparison with other tools

Results were evaluated for the two types of transfers that GrAnnoT can perform: from genome to graph and from genome to genome. In both cases, the transfers were performed with the different tools described before when possible. Then, for each transferred feature, its positions provided by the different tools were compared. Given a feature, we consider two transfers as different if they placed the feature at different positions. A transfer is specific to a tool if it is different from all the other transfers. By definition, a feature transfer is also specific to a tool if the feature is only transferred by this tool.

We tested GrAnnoT, GraphAligner, VG *inject* and VG *annotate* for the transfer of the annotation of the cv Nipponbare (Kawahara et al., 2013b) to the rice pangenome graph. We tested GrAnnoT, Liftoff, VG *inject/surject* and ODGI *position* for the transfer of the annotation of the cv Nipponbare to the cv Azucena.

##### Genome to graph transfer

For the genome to graph transfer, only the gene features were used. Indeed, VG *annotate* requires the GFF3 features to have a *ID* and a *Name* attribute to be transferred, while GrAnnot does not. It was the case only for the gene and mRNA features in the IRGSP annotation of the cv Nipponbare. For simplicity, we thus only selected the gene features in this annotation, as the mRNA are always included in a gene, and did not add other type of features to test the transfers.

The three methods that do not perform alignment (GrAnnoT, VG *inject* and VG *annotate*) have the exact same results for all the features. For ~32% of the features transferred by GraphAligner (17,870 features out of 55,798), the output is different from the other tools (Figure 2a). However, when allowing a difference of 1 bp on the position on the path, ~88% of the GraphAligner-specific transfers (15,673 out of 17,870 transfers) are then considered identical to the transfers from the other tools (Figure 2b). Further verification showed that these 1 bp differences from GraphAligner are alignment errors, where 1 bp is missing in 5’ or 3’ in the transferred feature sequence. Such differences are minor and acceptable for certain applications, but not in the context of annotation. Because of that, the current version of GraphAligner does not seem to be suitable for precise annotation transfer.

**Figure 2.**
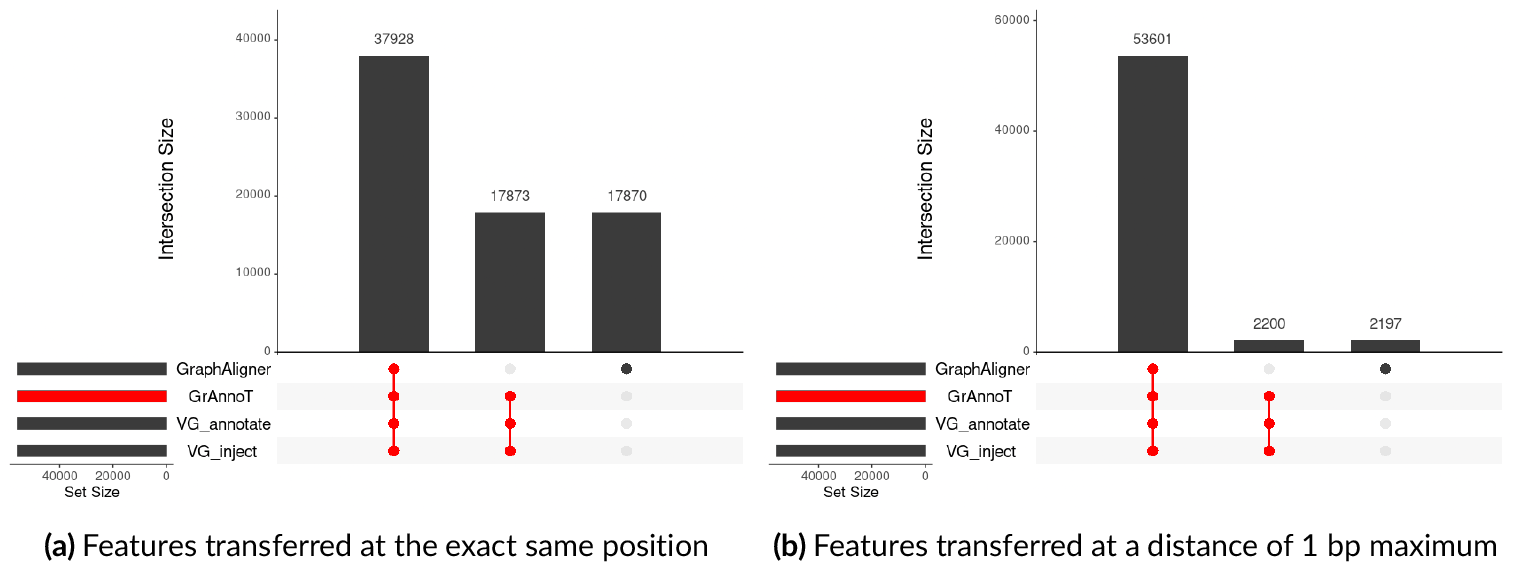
Genome to graph transfer comparison, Upset representation. Each vertical bar represents the number of identical transfers between the different tools specified below the bar. Two transfers are considered identical if they placed the feature at the exact same path in the graph and either at the exact same position on the nodes (a) or at a distance of maximum 1 nucleotide (b). The horizontal bars on the left represent the total number of transfers for each tool. GrAnnoT transfers are highlighted in red.

##### Genome to genome transfer

Most of the transfers between genomes are identical between the four tools (663,665/918,973≈72%). GrAnnoT seems to be the most consensual tool as it has the least specific transfers (Figure 3a) compared to the other tools. Liftoff has the most transfers, and seems to perform better than other tools (see below).

**Figure 3.**
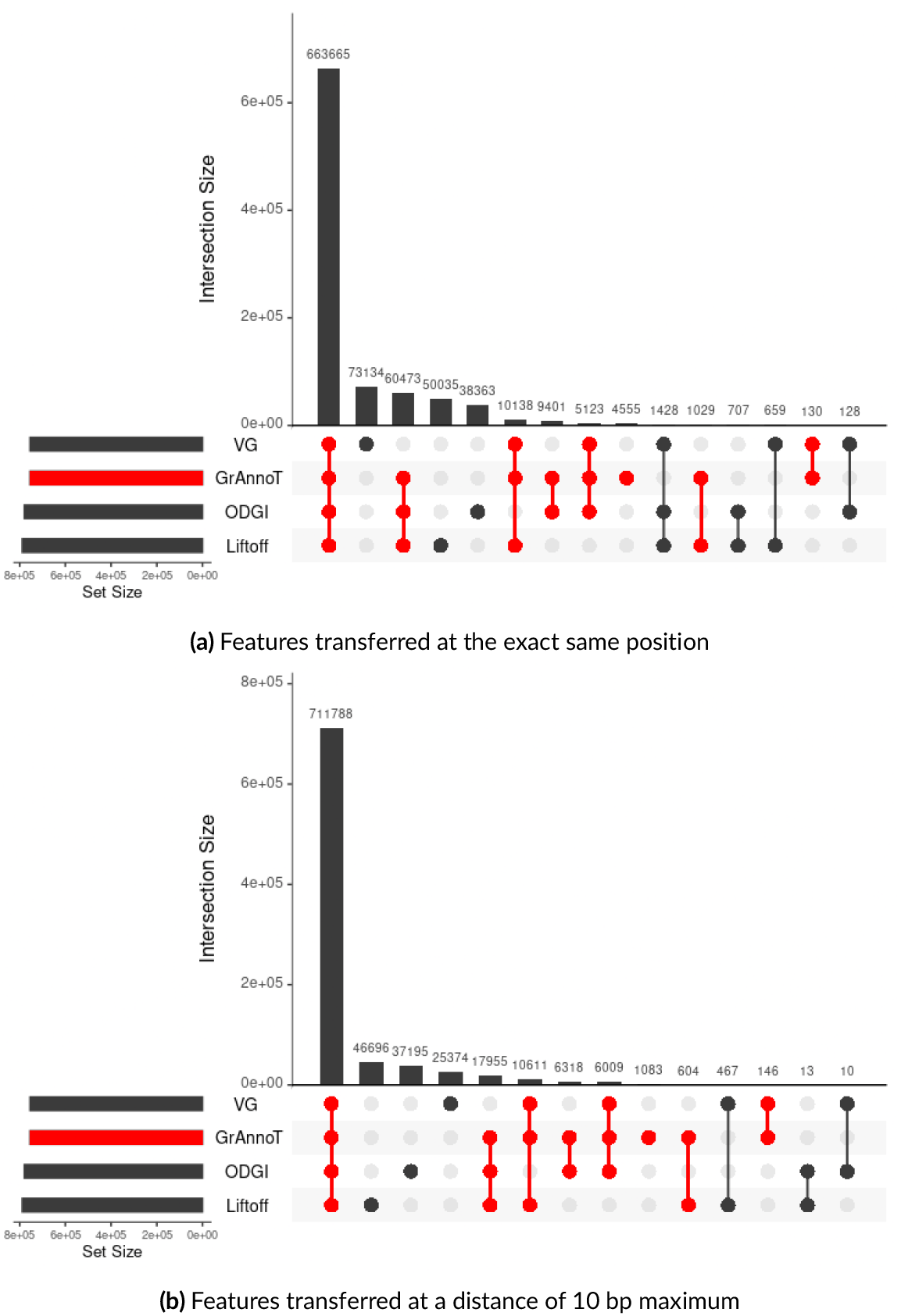
Genome to genome transfer comparison, Upset representation. Each vertical bar represents the number of identical transfers between the different tools specified below the bar. Two transfers are considered identical if they placed the feature either at the exact same positions on the target genome (a) or at a distance of maximum 10 nucleotide (b). The horizontal bars on the left represent the total number of transfers for each tool. GrAnnoT transfers are highlighted in red.

When looking at the tool-specific transfers, VG stands out the most, with 73,134 specific transfers. However, when allowing a difference of 10 bp between the transfers, VG has ~65.3% less specific transfers. Some of these VG specific transfers were manually compared to the transfers from the other tools for the same feature, and were identified as errors from VG (see supplementary Figure S10 for an example). This suggests that VG produces small errors during the *surject* step, since VG *inject* was shown to have the exact same results as GrAnnoT in Figure 2a. The 10bp difference tolerance revealed Liftoff and ODGI as the most divergent tools (with 46,696 and 37,195 specific transfers, respectively; Figure 3b).

Regarding the Liftoff-specific transfers, most are features that only Liftoff can transfer. Indeed, ~56% of them are inter-chromosomal translocations, *i*.*e*. features that are on a different chromosome between the source and the target genome (Figure 4). These transfers cannot be performed with GrAnnoT, VG or ODGI, as variation graphs are currently built chromosome-per-chromosome to reduce complexity, and therefore cannot represent such events (Andreace et al., 2023; Mergez et al., 2024). Thus, features on different chromosomes between Nipponbare and Azucena cannot be transferred by any of the graph-based approaches, and are found only by Liftoff. This could explain why Liftoff has the most transfers between the four tools.

**Figure 4.**
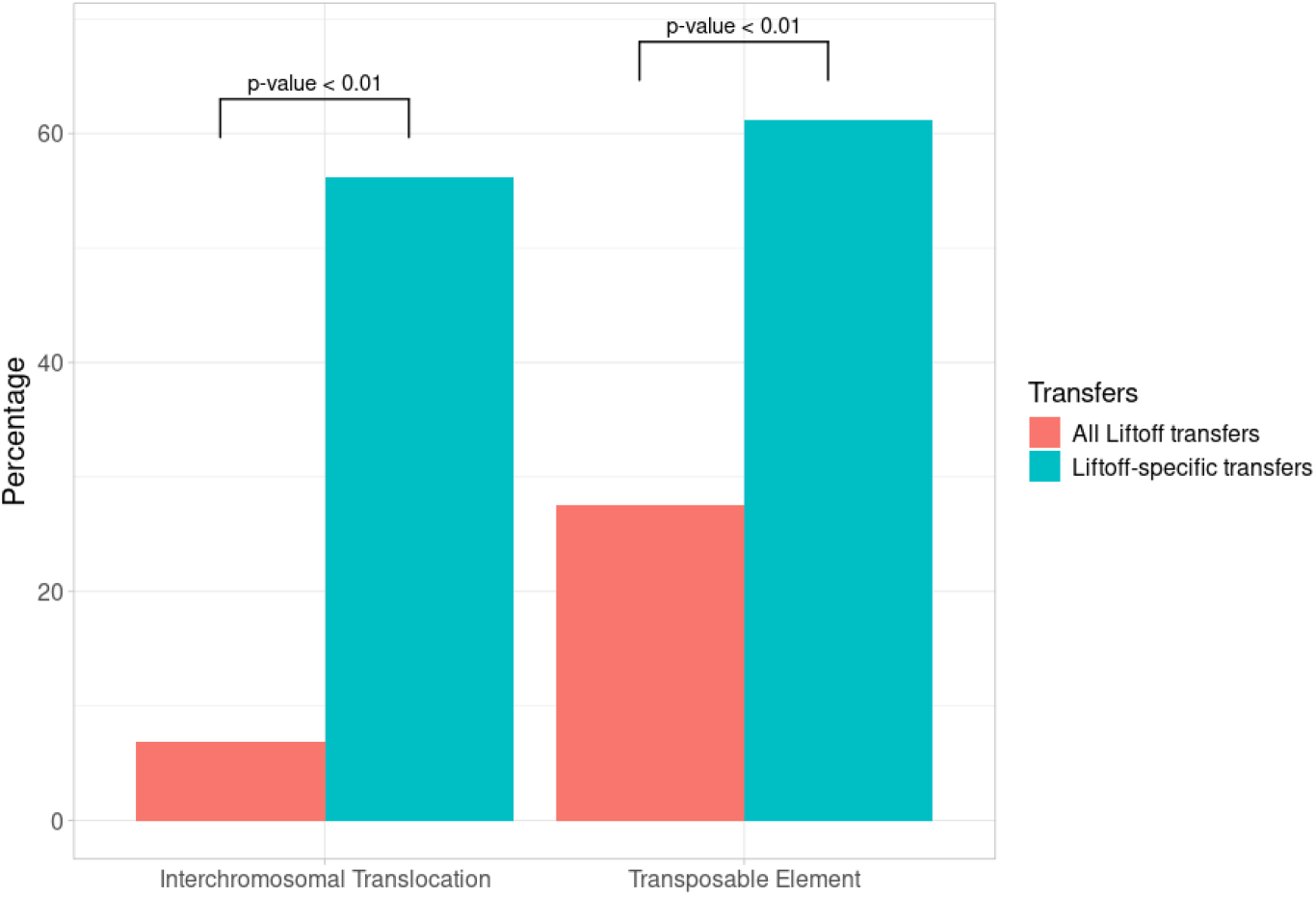
Transposable element and inter-chromosomal translocation percentages in all Liftoff transfer *vs* Liftoff-specific transfers. The Liftoff-specific transfers are enriched in interchromosomal translocations and transposable elements compared to all the other Liftoff transfers. Detailed data and *p*-value calculation are available in supplementary data (Tables S6 and S7).

Furthermore, when the annotations of these Liftoff specific-transferred features were thoroughly looked at, it appeared that they are enriched in transposable elements (TE) (*p*-value < 0.01; Figure 4). This could explain this discrepancy of chromosomal location between the two varieties, since transposable elements are mobile in the genome and can jump between chromosomes (Hayward and Gilbert, 2022; Wicker et al., 2007). The Liftoff-specific transfers that are on the same chromosome are also enriched in TEs (*p*-value < 0.01; Figure 5), as their ability to move in the genome makes them often not syntenic: encoding the relationships between such TEs in the graph with the current variation graph tools still seems complex (Eizenga et al., 2020). Indeed, the graph we used was built by minigraph-cactus, that aligns genomes to the graph in construction (Li et al., 2020). During this process, a non-syntenic region is more difficult to align, and its sequence can be represented in the variation graph by two different nodes carrying the same information. Because of that, some duplications, inversions, translocations, and TEs are not detected in the graph (Lemaitre, 2021; Romain et al., 2025), and annotations in these regions are not transferrable using only the variation graph structure. Since the relationships between the TEs are not always correctly encoded by the graph, TE annotation transfers cannot be reliably performed by tools such as GrAnnoT, VG or ODGI, that only use the structure of the graph.

**Figure 5.**
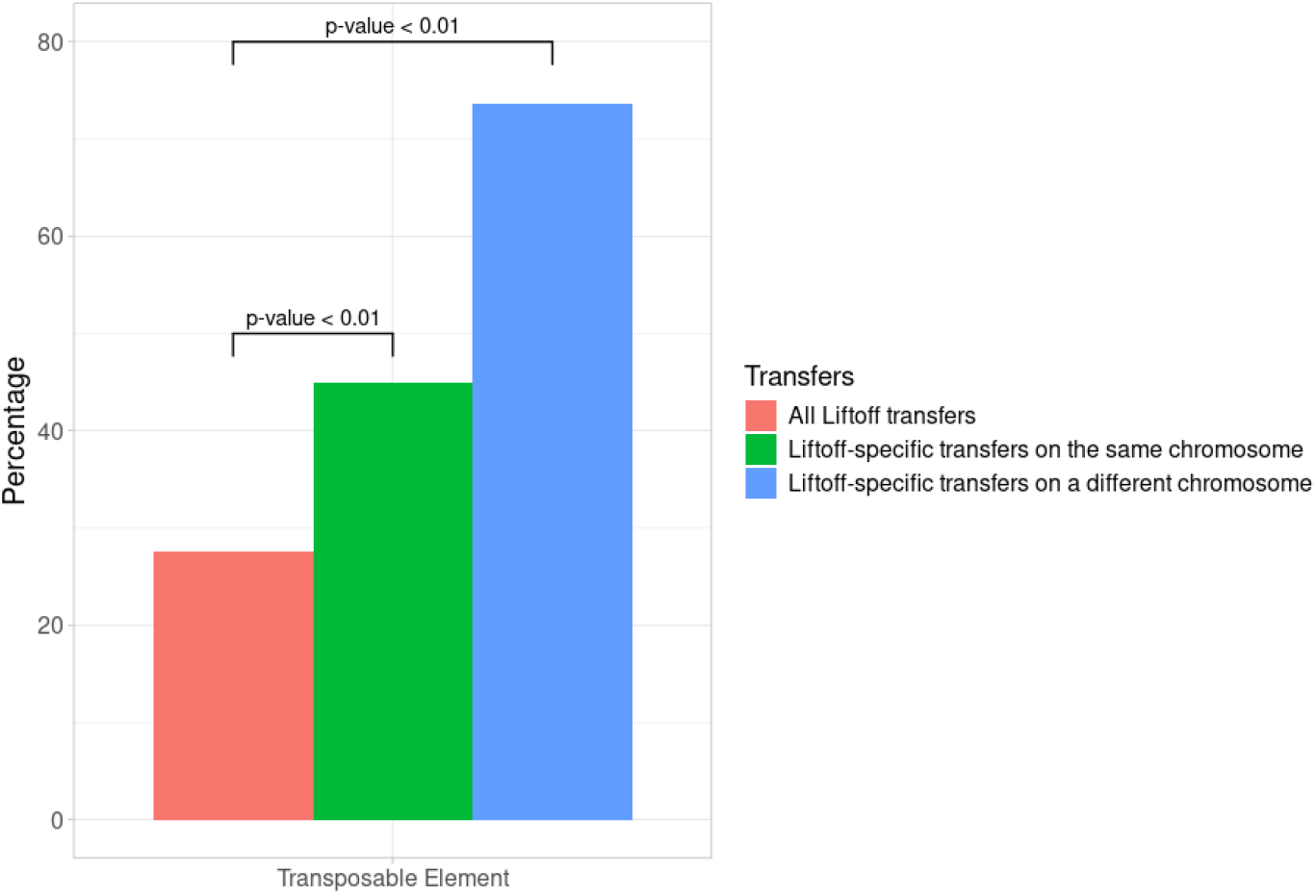
Transposable element percentages in all Liftoff transfer *vs* Liftoff-specific transfers. Compared to the other Liftoff transfers, Liftoff-specific transfers are enriched in transposable elements, whether or not they are in a translocation. Detailed data and *p*-value calculation are available in supplementary data (Tables S8 and S9).

Overall, Liftoff-specific transfers seem valid, and demonstrate a limitation in the variation graph approaches: they completely rely on the variation graph structure, which is not perfect and struggles to connect non-syntenic shared elements. The three graph tools tested are thus not suited for studying mobile sequences, such as interchromosomal translocations or TEs.

Most of the ODGI-specific transfers place a feature on a very small interval on the target genome. For instance, among the 37,195 ODGI-specific transfers, ~65% of the features (24,058) are placed on an interval of a length of zero nucleotide, and ~30% (11,320) on an interval of a length of one nucleotide. These transfers should be discarded, as they are of no biological meaning in terms of genes.

#### Robustness

##### Back and forth transfer

Two consecutive transfers (back and forth) with Liftoff and GrAnnoT allowed to compare how conservative these tools are. The first transfer was performed from the cv Nipponbare to the cv Azucena with the two tools. Then, the resulting Azucena GFF3 file was used as source to perform the second transfer, from the cv Azucena back to the cv Nipponbare. The resulting GFF3 for the cv Nipponbare was compared to its first original annotation in order to measure the loss or corruption of information during these transfers.

Liftoff loses less features during the two-round process (Table 1). This can be explained by the fact that Liftoff is better at finding non-syntenic features and handles interchromosomal translocations, as shown previously. However, while GrAnnoT did not lose any more annotation on the way back to the original sequence, Liftoff lost 257 features in the way back. In addition, when comparing the positions of the features before and after the two transfers (original *vs* transferred twice), GrAnnoT shows better results that Liftoff, with only 3.8% of the features being located at a different position compared to the original annotation, *versus* 10.4% of discrepancies for Liftoff. Moreover, after manual verification, it appeared that all the features misplaced by GrAnnoT in the second transfer are features where an extremity was shortened during the first transfer due to a deletion at the beginning or the end of the feature. Thus, while the feature transferred during the second transfer was incomplete regarding the true annotation, the transfer itself occurred correctly.

**Table 1.** GrAnnoT and Liftoff comparison on back and forth transfer. The input annotation for first transfer included 55,986 features. The loss corresponds to the number of features not transferred in either transfer (first cv Nipponbare to cv Azucena or second cv Azucena to cv Nipponbare). The other rows show how many features were at the same or at different positions before and after the two transfers.

|  | GrAnnoT | Liftoff |
| --- | --- | --- |
| Loss in first transfer | 7,961 | 1,622 |
| Loss in second transfer | 0 | 257 |
| Total loss | 7,961 | 1,879 |
| Same position | 47,184 | 48,482 |
| Different position | 841 | 5,625 |
| 1-10bp difference | 393 | 360 |
| 11-100bp difference | 275 | 648 |
| 101-1000bp difference | 165 | 776 |
| >1000bp difference | 8 | 1,210 |
| Different chromosome | 0 | 2,631 |
| Total transfers | 48,025 | 54,107 |

Finally, some features transferred by Liftoff back to the cv Nipponbare are placed on a different chromosome than the original one, as the transfer is alignment-based only and does not rely on synteny. In this regard GrAnnoT is more conservative than Liftoff. Indeed, orthologous copies are sometimes considered to guarantee a better conservation of gene function compared to paralogous copies, according to the ortholog conjecture (Nevers et al., 2020; Rogozin et al., 2014). As the variation graph conserves the synteny, GrAnnoT is more likely to transfer annotations between orthologous copies than between paralogous copies.

Overall, while GrAnnoT transfers less annotations than Liftoff (as seen in Figure 3a), it is more conservative with the annotations it does transfer.

##### Impact of the reference genome for graph construction

The variation graphs used were built with minigraph-cactus, which requires a reference genome as anchor, that can thus bias the graph structure (Andreace et al., 2023). To test the replicability of the GrAnnoT approach, transfers through two different graphs were compared. The two graphs have the same genomes embedded (13 Asian rice genomes), but were built with a different reference genome to initiate the graph. The reference genomes used for the two graphs are the annotated genome IRGSP-1.0 (cv Nipponbare), and Os127652RS1 (cv Natel Boro) (Zhou et al., 2020). Annotation transfer from cv Nipponbare to cv AzucenaRS1 was performed with these two graphs, and the positions of the common transferred features were compared.

Among the 48,322 features transferred on the cv Azucena, 2,673 (~5.5%) were not transferred by both graphs. Among the 45,649 features transferred by both graphs, only 393 (~0.9%) were not transferred at the same exact location (Table 2).

**Table 2.** Comparison of GrAnnoT transfers using graphs with different reference genomes. The input annotation for the transfer included 55,986 features. The loss corresponds to the number of features not transferred. The other rows show how many features were placed at the same or at different positions when transferred with the two graphs.

|  | Nipponbare reference | Natel Boro reference |
| --- | --- | --- |
| Total transfers | 48,025 | 45,946 |
| Loss | 7,961 | 10,040 |
| Specific transfers | 2,376 | 297 |
| Comparison between the two graphs |  |  |
| Common transfers | 45,256 |  |
| Different transfers | 393 |  |
| 1-10bp difference | 169 |  |
| 11-100bp difference | 87 |  |
| 101-1000bp difference | 76 |  |
| >1000bp difference | 61 |  |
| Different chromosome | 0 |  |

The amount of features not transferred by both graphs is not negligible, even though they mostly consist of TEs (see before). However, it can be explained by the choice of the reference genome for the second graph construction, the cv Natel Boro. Indeed, among the 11 genomes in the graph that are not involved in the transfer (neither the cv Nipponbare nor the cv Azucena), the cv Natel Boro is among the furthest, genetically speaking, as shown in the phylogenetic tree in the genomes original paper (Zhou et al., 2020). Thus, it makes sense that the graph centered around the cv Nipponbare displays better performance for annotation transfer from the cv Nipponbare. This showcases the importance of the choice of the reference genome for the graph construction, that must be adapted to the use case of the graph.

##### Comparison with other species

GrAnnoT was compared to Liftoff using two other datasets: a pangenome variation graph of the human chromosome 1 (Liao et al., 2023) and an *E. coli* pangenome variation graph (**data_ecoli**), both made with minigraph-cactus. For the rice graph, the transfer was again made from the cv Nipponbare to the cv Azucena; for the human graph, the transfer was made from the CHM13 to the GrCH38 haplotype; for the *E*.*coli* graph, the transfer was made from the O157_H7_EC4115_0a2c271 strain to the S88_fa4fe08 one. These comparisons checked if the positions of the features transferred by both approaches are consistent, to assess if the results observed in the rice pangenome graph were replicable with graphs from other type of dataset/organisms/phylum.

It appears that for the rice and human datasets, most of the features are transferred by both tools (~85.7% for rice and ~91.3% for human) (Table 3). For *E. coli*, only ~62.5% of features are transferred by Liftoff and GrAnnoT, and both tools have relatively low transfer rates (below 70%, see Table 3). This suggests that the features not transferred by either tool reflect a difference in gene content between the two strains, rather that a technical error.

**Table 3.** Comparison between GrAnnoT and Liftoff in several species. Each feature in the input annotation was transferred using GrAnnoT and Liftoff. When the feature has been transferred by both tools, the two positions given were compared to see how different they are.

|  | Rice |  | Human Chr1 |  | E.coli |  |
| --- | --- | --- | --- | --- | --- | --- |
| Total features to transfer | 55,986 |  | 282,668 |  | 11,460 |  |
| Features transferred by GrAnnoT | 48,025 | 85.78% | 276,681 | 97.88% | 7,407 | 64.63% |
| GrAnnoT-specific transfers | 72 | 0.13% | 4,647 | 1.64% | 239 | 2.09% |
| Features transferred by Liftoff | 54,363 | 97.10% | 275,249 | 97.38% | 7,939 | 69.28% |
| Liftoff-specific transfers | 6,410 | 11.45% | 3,264 | 1.12% | 767 | 6.69% |
| Features transferred by both tools | 47,951 | 85.65% | 258,092 | 91.31% | 7,167 | 62.54% |
| Same position | 46,431 | 82.93% | 256,927 | 90.89% | 6,844 | 59.72% |
| Different position | 1,520 | 2.71% | 1,165 | 0.41% | 323 | 2.82% |
| 1-10bp difference | 795 | 1.42% | 623 | 0.22% | 184 | 1.61% |
| 11-100bp difference | 284 | 0.51% | 150 | 0.05% | 70 | 0.61% |
| 101-1000bp difference | 155 | 0.28% | 46 | 0.02% | 15 | 0.13% |
| >1000bp difference | 164 | 0.29% | 346 | 0.12% | 54 | 0.47% |
| Different chromosome | 122 | 0.22% | 0 | 0% | 0 | 0% |
| Runtime Liftoff | 00:23:45 |  | 00:10:27 |  | 00:00:09 |  |
| Runtime GrAnnoT | 00:08:11 |  | 00:14:47 |  | 00:00:22 |  |

Additionally, a large part of the features transferred by both tools are placed at the exact same position by Liftoff and GrAnnoT (~96.8% for rice, ~99.6% for human and ~95.5% for *E. coli*). As expected, some features are transferred only by Liftoff, but for the human graph GrAnnoT-specific transfers appear in negligible quantities. This better transfer capacity for the two tools in human may be due to the lesser diversity of human genomes compared to rice (mean 15.6 millions SNP for 64 human haplotypes *vs* 9.4 millions for only 16 rice ones, respectively; Ebert et al., 2021; Wei et al., 2024), and even more so compared to *E. coli*. In addition, the annotation of human genes is probably better curated than in rice, with less hypothetical genes that may be false positive, also explaining the better transfer for both tools on human reference. Liftoff-specific transfers for E.Coli were manually inspected and most of them are related to unknown protein domains or related to biotic and abiotic stress responses. Such type of genes in bacteria are generally related to mobile structures such as ICE, e.g. Zheng et al., 2023.

##### Scalability

To ensure GrAnnoT can work with larger datasets, it was tested on a variation graph build using minigraph-cactus with 69 *A*.*thaliana* genomes (Lian et al., 2024; Mergez et al., 2024). GrAnnoT was used to transfer the annotation of the genome *Abd-0* to the graph and then to all 68 other genomes. This operation was performed in ~2h52min, with an average of ~2min30sec per transfer (including the transfer from *Abd-0* to the graph).

### 2.3. Run time comparison

The execution time for the transfer from genome to genome with the different tools was measured on Asian rice data between the cv Nipponbare and the cv Azucena, using the command */usr/bin/time* (Table 4). The graph tools use as input the annotation file in GFF3 and the variation graph in the adapted format. VG *inject/surject* (as we deal here with genome to genome transfer) and ODGI *position* require the variation graph to be converted/indexed in their format before use (.xg and.og, respectively), and GrAnnoT uses directly the variation graph in native GFA format. Liftoff uses as input the fasta files for the genomes and the annotation file in GFF3 format. The results show that GrAnnoT has the best run time, and that ODGI and VG *inject/surject* are substantially slower that GrAnnoT and Liftoff.

**Table 4.** Run time comparison for genome to genome transfer in rice. GrAnnoT, VG *inject/surject* and ODGI *position* transfer through the graph, and VG inject/surject and ODGI convert/index the graph before the transfer. Liftoff directly uses the genome fasta files.

|  | GrAnnoT | Liftoff | VG *inject/surject* | ODGI *position* |
| --- | --- | --- | --- | --- |
| Graph construction | 04:23:22 | - | 04:23:22 | 04:23:22 |
| Graph conversion/index | - | - | 00:14:34 | 00:05:37 |
| **Annotation transfer** | **00:08:11** | **00:23:45** | **07:12:17** | **70:18:04** |
| Total time | 04:31:33 | 00:23:45 | 11:49:73 | 74:46:63 |

GrAnnoT was further compared to Liftoff in terms of run time.

Several transfers were performed with both tools to compare the run times, because GrAnnoT is designed to facilitate the transfer toward multiple target genomes: it starts by pre-processing the variation graph and loading the graph annotation, which only needs to be done once, no matter how many target genomes are included in the graph.

Liftoff can be run in GFF mode or in database mode; the database mode needs less time since the GFF annotation file has already been processed. Both of these mode were compared to GrAnnoT.

The commands timed are transfers from the cv Nipponbare to all the other genomes in the rice pangenome graph, with 12 transfers in total.

The results show that GrAnnoT is faster than Liftoff to perform one annotation transfer (~8 minutes *vs* ~22 minutes), and even more to perform twelve (~47 minutes *vs* ~5 hours and 30 minutes, see Figure 6). However, this comparison does not take into account the time needed to build the graph. When adding the graph construction time (~4h23mn on our infrastructure) to the GrAnnoT 12 transfers time, we still get a duration (~5h10mn) slightly shorter to Liftoff transfers (~5h47min or ~5h29min). Additionally, GrAnnoT can give supplementary informative output that describe the transfers performed such as a presence-absence matrix or alignment files of the transferred features, as well as the graph annotation.

**Figure 6.**
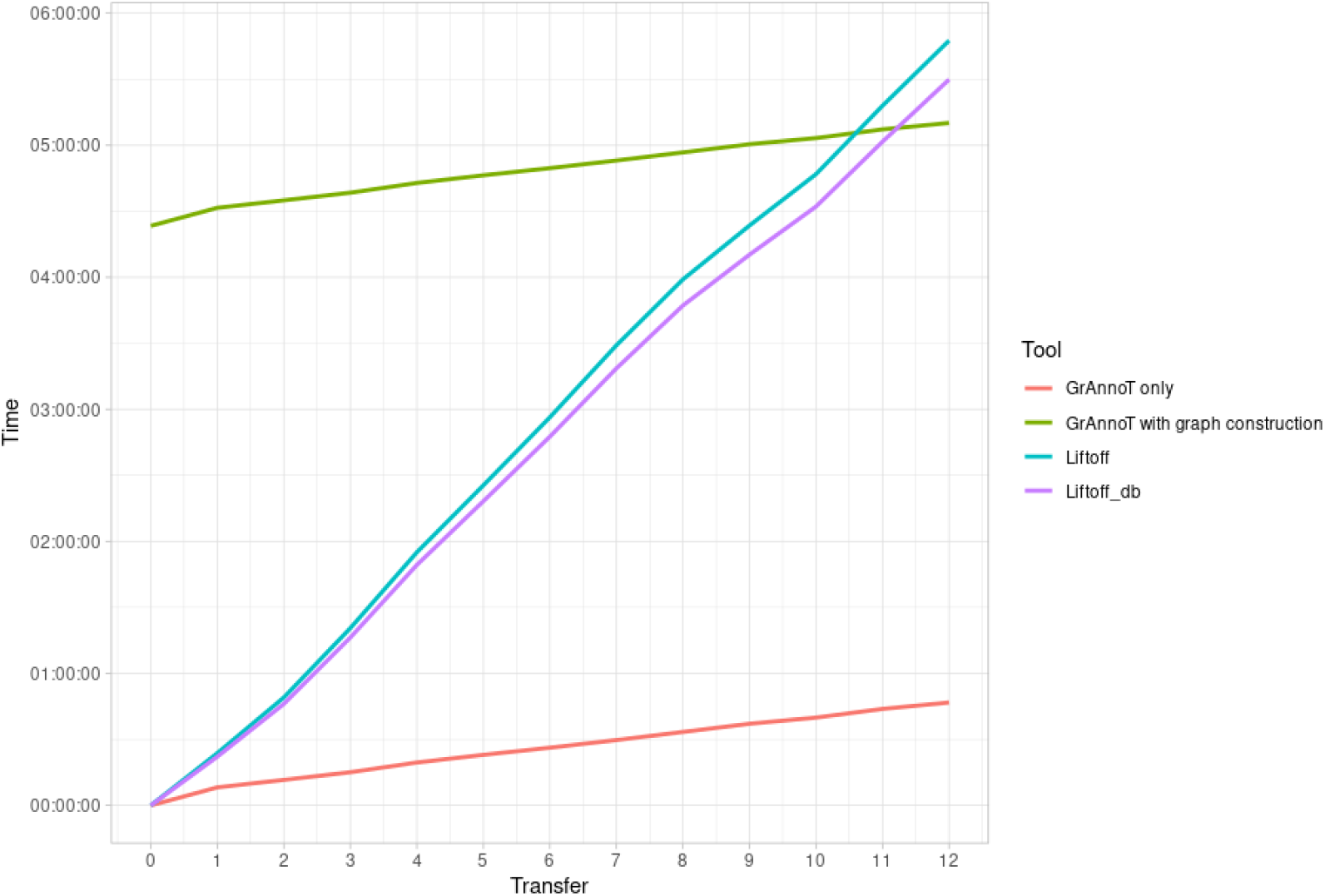
Genome to genome transfer comparison. GrAnnoT and Liftoff run time for 1-12 transfers were measured using the command */usr/bin/time*. Liftoff was run both in GFF and DB mode. GrAnnoT values are presented including or not the graph building time. Detailed time points are available in supplementary data in Table S10.

For the human graph, GrAnnoT is not faster than Liftoff for one transfer (see the last lines of Table 3). However, as shown on Figure 6, for several transfers GrAnnoT is more advantageous. We tested the runtime of GrAnnoT for the annotation transfer on 10 haplotypes, and got ~45 minutes in total. This is significantly lower than the time for one transfer multiplied by 10 (~1h44min), which is what we can expect of 10 Liftoff transfers from the results in Figure 6.

## 3. Applications

To assess the use of GrAnnoT annotation transfer, in particular the informative outputs complementary to the GFF3 itself, we analyzed a few characteristics of the annotation transfers between the Nipponbare and Azucena cultivars. More precisely, we verified that the variations in the graph reported by GrAnnoT are distributed as biologically expected, in a way that does not disrupt the proteins coded by the gene features.

### 3.1. Indel rate in different feature types

We looked at the positions of the indel variants (insertion or deletion) in the different feature types that correspond to different parts of the genes. These variants are expected to be less present in the CDS compared to the rest of the gene due to selection pressure, because the resulting changes in the coded protein are more important.

The feature types that were compared are:

- the whole gene feature itself
- the mRNA
- the 5’UTR
- the exons
- the CDS
- the introns
- the 3’UTR

These feature types have different average lengths, inducing a bias in the number of indel found by feature type; if the indels are randomly distributed, we expect more indels in the feature type that has the longest cumulated length. To counter this bias, for each feature type we reported the total number of indels found to its cumulated length, obtaining the average number of indel per position.

The results displayed in Figure 7 show that, as expected, the CDS have the fewest indels and the non-coding regions (UTR and introns) have the most. This confirms that the variations in the graph reported by GrAnnoT are consistent with the current understanding of genome variation selection.

**Figure 7.**
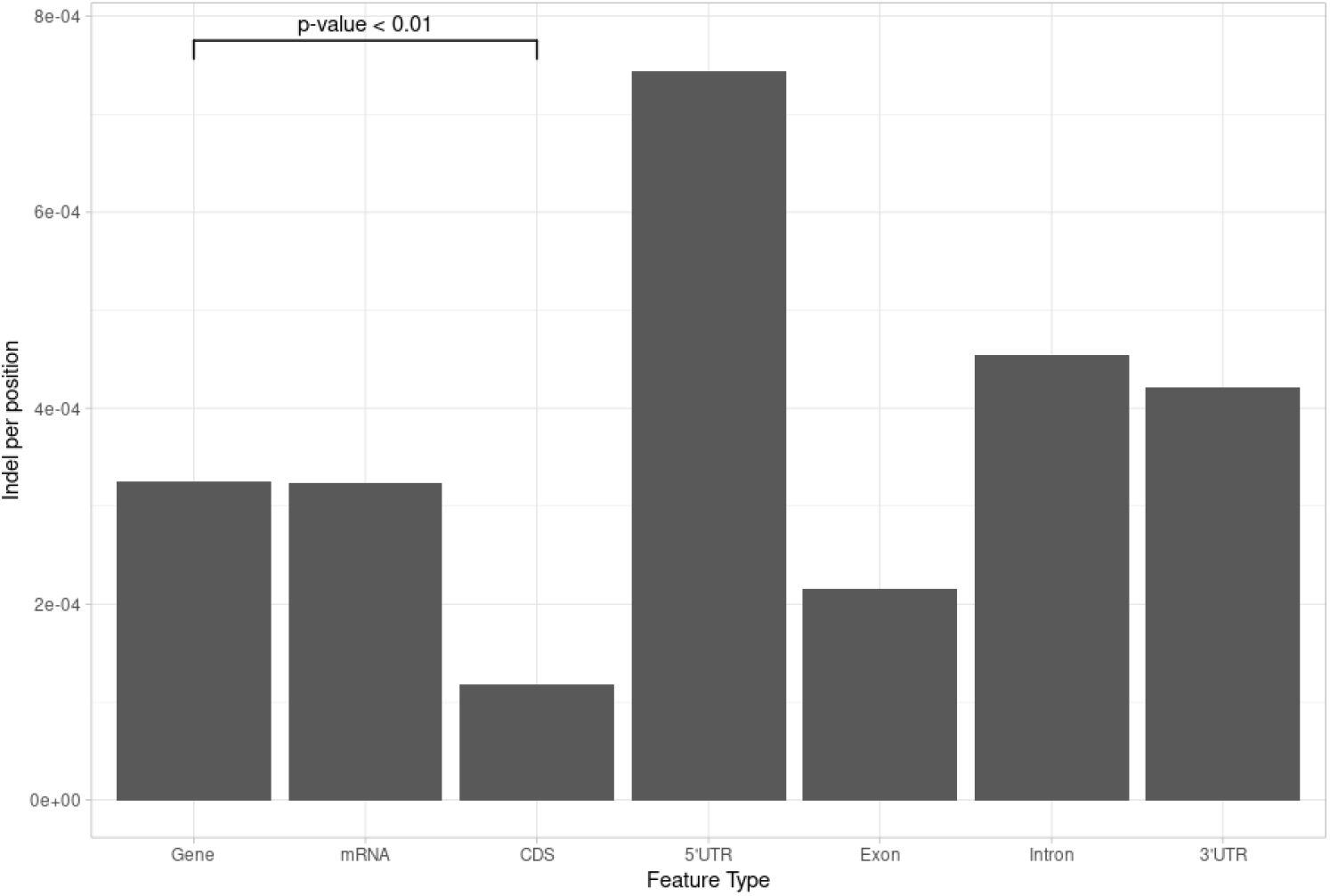
Indel count. Each bar represents the number of indels (insertion or deletion) per position in the corresponding feature type. As expected, the CDS are the most conserved and thus have the least indels, and the non-coding regions (UTR and introns) are the least conserved and have the most indels. Detailed data and *p*-value calculation are available in supplementary data (table S11).

### 3.2. Frameshift mutations in different feature types

Indels can modify the protein coded by a gene, but indels in CDS are particularly impactful when they change the reading frame. We calculated the rate of frameshift mutations (indel whose length is not a multiple of 3) among the indels, for all feature types. We expect to have a lower ratio of frameshift mutations in the CDS compared to the non-coding regions, because of the selection pressure.

The results displayed in Figure 8 show that the CDS have the lowest percentage of frameshift variation from their indels, and that the introns have the highest.

**Figure 8.**
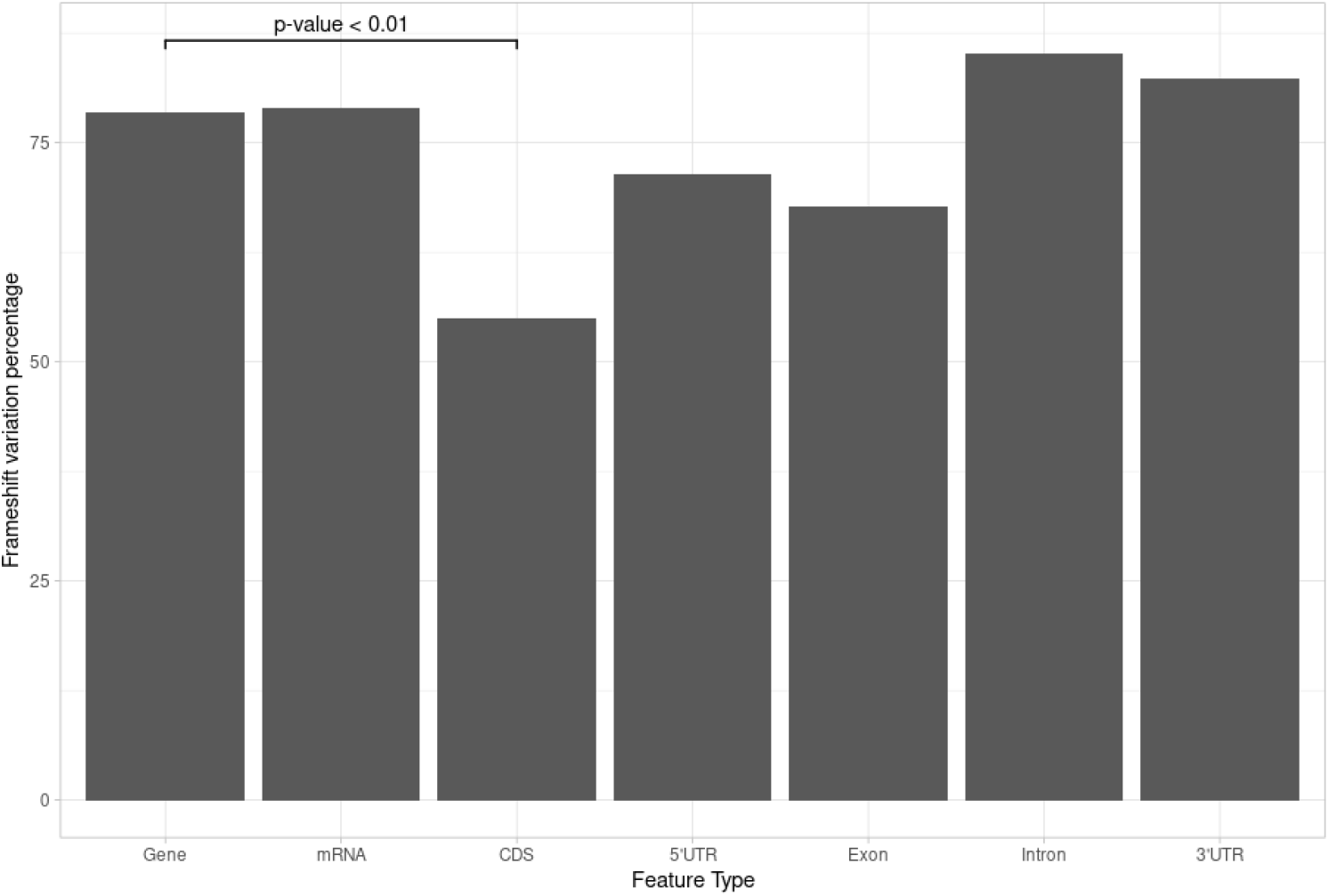
Frameshift indel count. Each bar represents the percentage of frameshift variant (length not multiple of 3) among all the indels in each feature type. As expected, the CDS have the fewest frameshift variant, since these variants impact significantly the protein coded. Detailed data and *p*-value calculation are available in supplementary data (table S12).

### 3.3. Substitutions position in different feature types

Substitutions are usually smaller variants than indels, so they are expected to have a smaller impact. However their distribution in CDS is not expected to be uniform. Indeed, substitutions on the third position of a codon is more likely to be silent than a substitution on the two other positions. Because of that, in CDS the third codon position usually has more substitutions than the two other positions (Sanchez et al., 2005).

On Figure 9, we show that the CDS indeed has more substitutions on the third codon position than the other two positions, while the other gene elements have more homogeneous substitution distributions. Details about how we computed the substitution position in the CDS are available in supplementary data (3).

**Figure 9.**
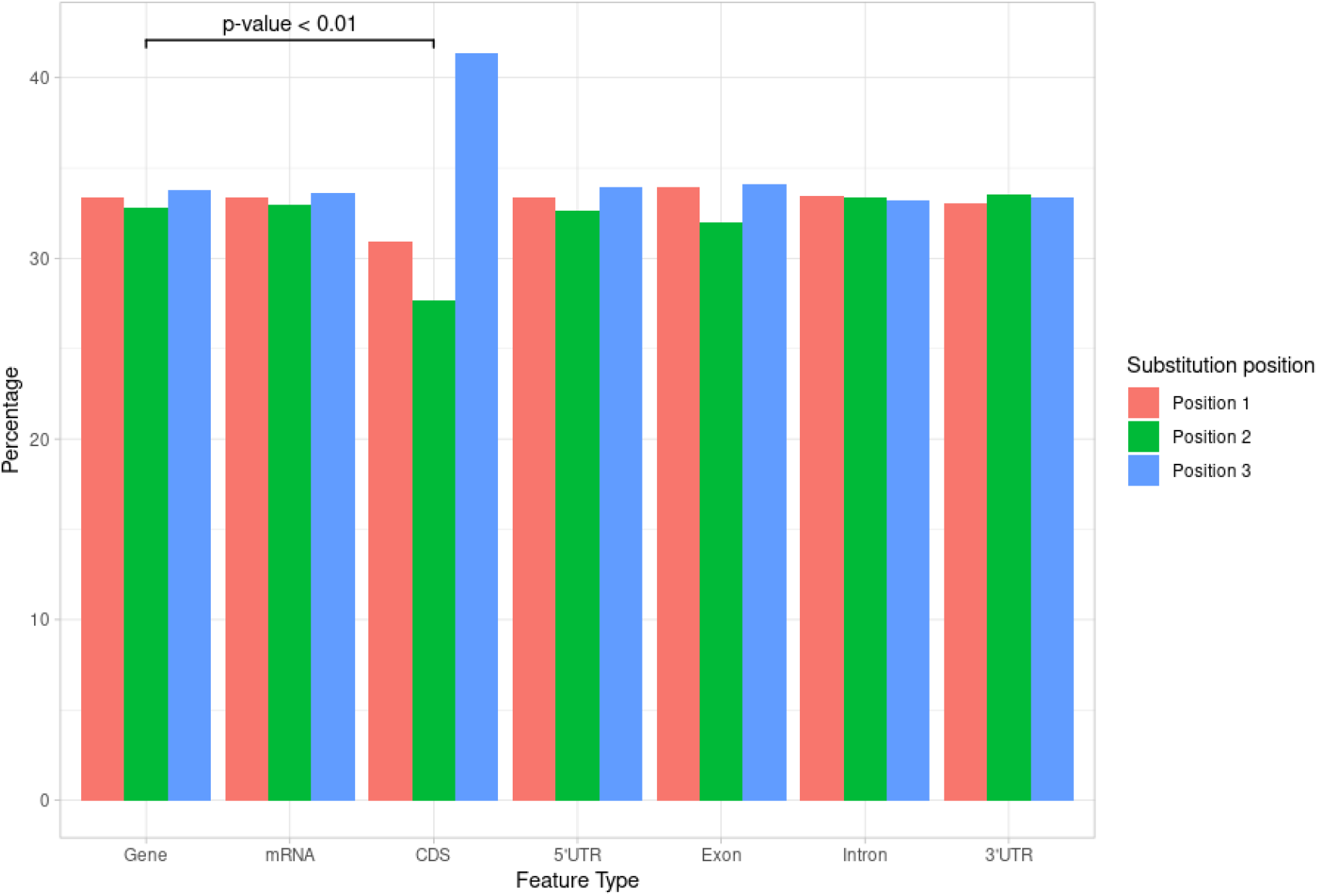
Substitution positions. For each feature type, the percentage of substitutions that are on each of the three codon positions is displayed. In the CDS, the third position has more substitutions than the two other positions. For the other feature types, we don’t see that the positions multiple of 3 have more substitutions than the others. Detailed data and *p*-value calculation are available in supplementary data (table S13).

### 3.4. Pangene set analysis

The PAV matrix output was computed for all the genomes in the Asian rice graph (minus the source genome cv Nipponbare), and was used to compute the core, dispensable and shell gene set from the cv Nipponbare in this pangenome.

We found ~58% of core genes and ~33% of dispensable genes (table 5) in our variation graph, which is similar to what is seen in the literature when accounting for the different threshold chosen in each study (with ~53-62% of core and ~38% of dispensable gene families for instance; Wang et al., 2018).

**Table 5.**
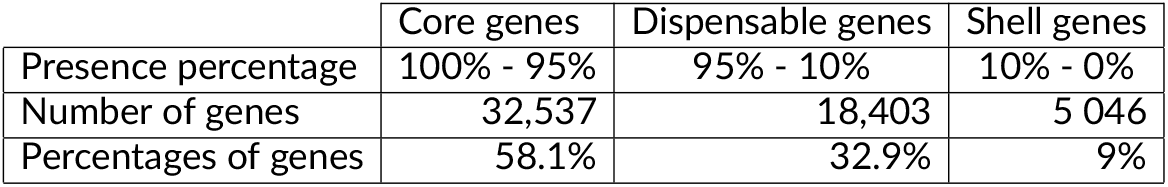
Core, dispensable and shell gene set. The population size is 12, and there are 55,986 genes in total.

## Discussion and conclusion

Annotation of pangenomes is a broad and crucial topic, that has been explored in various ways depending on the type of pangenome considered. Some pangenomes consist in a gene set, where several genomes are separately annotated and the resulting genes are clustered into orthologous groups (Gautreau et al., 2021; Gordon et al., 2017). Alternatively, pangenomes can be stored in de Bruijn graphs, that ggCaller (Horsfield et al., 2023) can annotate *de novo* (for bacteria), or that Pantools (Jonkheer et al., 2022) can visualize and project annotation on this visualization. However the variation graph structure has less manipulation options, and most of them rely on the visualization on the graph (Liu et al., 2024; Miao and Yue, 2025). Visualizing whole variation graphs can be challenging, as these structures are usually complex and non-linear for large genomes, resulting in a hairball-like structure which is difficult to interpret (Durant, 2022). It is easier to project and visualize only a few annotations on a targeted region of the graph to study the variations, but this approach is less scalable, poorly reproducible, and usually requires to know in advance which regions to study.

GrAnnoT fills this gap by efficiently projecting an annotation on a variation graph and its embedded genomes, and providing useful information on the variants in the annotated regions between the different genomes embedded in this graph. It is easy to install and to use, and the operations it proposes are fast, reliable, reproducible, and it gives informative and user friendly outputs. Once the annotation has been transferred in the variation graph and its variations between the genomes have been reported, subgraphs of the regions or genes of interest can be extracted with tools like VG or ODGI, or GrAnnoT in the future. A visualization of these subgraph can help study in more details the variations found in the regions targeted with GrAnnoT.

The main limitation of GrAnnoT, as displayed in this paper, comes from the variation graph itself : GrAnnoT solely relies on this graph structure and does not perform any alignment (in order to conserve speed); thus, any flaw in the variation graph will hinder GrAnnoT in its transfers. As a first example, graphs built by minigraph-cactus (such as the ones we used) separate the chromosomes to build independent graphs, and thus cannot represent interchromosomal events (such as translocation or transposition). In addition, the example of the transposable elements in the Asian rice dataset highlights that minigraph-cactus currently struggles to align some non syntenic elements (transposed copies), even on the same chromosome. Therefore, those annotations cannot be transferred by GrAnnot. GrAnnoT can handle gene duplication and inversion when the information is present in the variation graph; however, the detection of these variants (by minigraph-cactus) can be inconsistent (Lemaitre, 2021; Romain et al., 2025). When the graph builders will be able to inform on interchromosomal relationships and duplications, GrAnnot will be able to immediately transfer the annotations corresponding to these regions.

Indeed, applying GrAnnoT to a graph built with another tool than minigraph-cactus would be an interesting option, to see if other graph builders are better at representing non-syntenic relationships.

To test that, GrAnnoT has been applied to PGGB graphs, as this graph builder has the benefit of being completely reference-free. The annotation transfer on the PGGB graph was performed correctly, but the transfer on a target genome was very slow and erroneous in repeated regions, for reasons not yet identified (data not shown). One possible explanation could be the tendency of PGGB graphs to have many cycles, at the opposite of minigraph-cactus graphs (Andreace et al., 2023). This resulting complexity could be the reason for the high runtime and errors of graph-to-genome transfers, as GrAnnoT algorithms are not suited for such complex graph topologies. However, in the future, we plan to improve the compatibility between PGGB and GrAnnoT, in order to be adaptable to more type of variation graphs.

Minigraph is another tool that builds pangenome graphs, but the output format (rGFA format) is incompatible with GrAnnoT as its graphs do not include paths or walks, and thus cannot inform to which path belong the current annotation.

GrAnnoT was compared to existing tools that perform some similar operations, but none correspond exactly to its scope or its full capacities.

For annotation transfer to the graph, GrAnnoT was compared with VG annotate, and the two tools gave the exact same results. Both are straightforward to use, and are good options. VG inject has the same behavior (outside of its complexity in use) as VG annotate, and thus is also efficient. On the opposite, GraphAligner is too slow and showed some errors in placing gene borders. The runtimes are in the same order of magnitude between GrAnnoT and VG annotate, with the VG strategy consisting in investing time to index the graph for rapid individual transfers later on, while GrAnnoT uses directly the GFA in its native form.

For annotation transfer between genomes, GrAnnoT has showed good results and is comparable to Liftoff in terms of performance for syntenic elements. As previously mentioned, Liftoff is better at transferring non syntenic annotations, and should thus be preferred by users interested in such elements. In terms of CPU time, once the graph is built GrAnnoT is faster than Liftoff, and the graph construction time can be compensated when performing several transfers. While GrAnnoT can transfer annotations between genomes as accurately as Liftoff, its primary objective is to integrate annotations to a pangenome variation graph, and therefore to inform the user of the variability of the population in terms of genes, and of the variation within the genes sequences. In comparison, Liftoff only provides a percentage of similarity, and not any alignment or information on the variants.

For performing both tasks, VG offers the commands *inject/surject*, and ODGI the command *position*. However the runtimes for these tools are prohibitive, and their transfers are not reliable, probably because they were aimed to transfer a few set of coordinates in non-complex, human genomic regions.

Another addition in GrAnnoT compared to VG and ODGI is the filtering based on sequence identity and coverage, which ensures that a substantial part of the annotated feature is present in the target genome. The threshold used can be easily modified by the user, to adapt to the species, pangenome diversity, graph quality, type of feature annotated, etc.

In the future, we plan to improve GrAnnoT capacity to annotate a pangenome variation graph by including more than one annotation. Indeed, GrAnnoT current approach for transferring annotation comes with the downside of only informing on regions that are shared with the originally annotated genome. However, one advantage of the pangenome is to study dispensable regions, that are not shared by every genome and thus that are not always present in the annotated genome. When available, integrating annotations from several genomes in the variation graph would give a more complete picture of the gene set in the population, and help study the whole pangenome.

Another future development of GrAnnoT could be to aim to reduce the loss of non-syntenic annotations. For that, we would first need to detect them, by selecting gene annotations poorly transferred for example. Then, using a graph alignment tool, we could find all the different sets of nodes representing their sequence, *i*.*e*. all the different locations of the annotated sequence in the graph. This approach would help transfer annotations lost due to transposition or chromosomal rearrangement, but requires accurate detection of elements not transferred because of the graph structure. Indeed, the alignment of gene sequences on the graph is time consuming, and naively aligning all the genes would be too long and redundant, as the graph correctly aligns most of the genomes regions.

In conclusion, the present study introduced GrAnnoT, the first tool able to efficiently transfer annotation on a pangenome variation graph from one of its embedded genomes and reverse. It relies on the already performed alignment that created the graph to identify syntenic segments. We benchmarked GrAnnoT on Asian rice, bacteria and human pangenomes, and showed that it is fast, scalable, reliable and efficient, and performs adequately compared to state-of-the-art tools for linear genomes. It is a robust, replicable tool working on any type of species for which a variation graph is available. In addition, GrAnnoT can provide useful outputs, such as the alignments of the gene sequence between source and target, or a presence/absence matrix.

## Acknowledgements

The authors want to thank Sebastien Ravel from PHIM/CIRAD for his valuable discussions and help on packaging and containerization.

The authors acknowledge the ISO 9001 certified IRD i-Trop HPC (South Green Platform) at IRD Montpellier for providing HPC resources that have contributed to the research results reported within this paper. URL: https://bioinfo.ird.fr/-http://www.southgreen.fr

## Data, script, code, and supplementary information availability

Data and results are available online (data: https://doi.org/10.23708/DO1RTF,Marthe and Sabot, 2025a, results: https://doi.org/10.23708/RRSKRA, Marthe and Sabot, 2025b). Scripts and codes are available online (https://doi.org/10.23708/TW3KYV, Marthe et al., 2025).

## Supplementary data

### Data and tools used

- Data
  - *E*.*coli* graph (built with data obtained using the same protocol as Heumos et al., 2024, see below)
  - Human graph (from Liao et al., 2023)
  - Rice graph (built with data from Kawahara et al., 2013a; Zhou et al., 2020)
- Tools
  - minigraph-cactus v2.8.2 (Hickey et al., 2023)
  - Liftoff v1.6.3 (Shumate and Salzberg, 2021)
  - ODGI v0.8.6-11-ga1f169cc (Guarracino et al., 2022)
  - VG v1.58.0 (Garrison et al., 2018)
  - GraphAligner Branch master commit daec67f67a2f50d648a6aa30cbbe5a2949583061 (Rautiainen and Marschall, 2020)

### NCBI ID of *E*.*coli* genomes used to build the graph

- NC_000913.3
- NC_002655.2
- NC_004431.1
- NC_007779.1
- NC_008253.1
- NC_008563.1
- NC_009800.1
- NC_010468.1
- NC_010473.1
- NC_011353.1
- NC_011601.1
- NC_011741.1
- NC_011742.1

### Substitution positions

To find the codon positions for the subsection 3.3, we had to take into account the splicing of the mRNA. Indeed, the CDS elements in the annotation only correspond to a fraction of the real CDS in the mRNA. Thus the substitution positions are not relative to the real CDS, and finding the third position of the codon required to add the context of the preceding CDS fragments. This adjustment was only done for the CDS elements in the annotation, since they are the only splited elements. This explains why the exons do not follow the CDS tendency in Figure 9, contrary to Figures 7 and 8.

**Figure S10.**
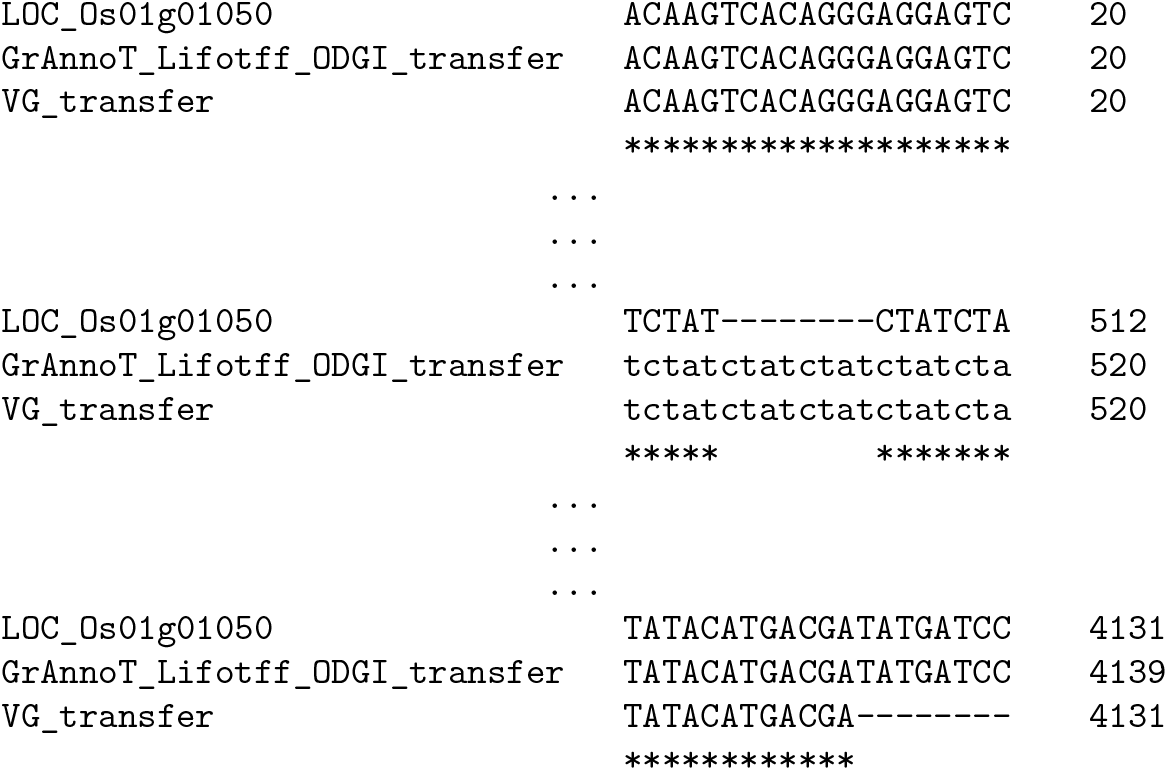
Extraction of the alignment of LOC_Os01g01050 gene and its transfers on cv Azucena by different tools. VG *inject/surject* transfer appears to have an error as the positions it gives miss the last 8 bases of the gene. The gene total length is conserved in VG transfer because there is an insertion in cv Azucena in the middle of the gene.

**Table S6.** Interchromosomal translocation rates in Liftoff transfers. The *p*-value measures the enrichment in interchromosomal translocations in the Liftoff-specific transfers, and was computed with Pearson’s Chi-squared test.

|  | Different chromosome | Same chromosome | *P*-value |
| --- | --- | --- | --- |
| Liftoff-specific transfers | 3,604 | 2,806 | < 0.01 |
| Other Liftoff transfers | 122 | 47,831 |  |

**Table S7.** Transposable elements rates in Liftoff transfers. The *p*-value measures the enrichment in transposable elements in the Liftoff-specific transfers, and was computed with Pearson’s Chi-squared test.

|  | Transposable elements | Other features | *P*-value |
| --- | --- | --- | --- |
| Liftoff-specific transfers | 3,919 | 2,491 | < 0.01 |
| Other Liftoff transfers | 11,053 | 36,900 |  |

**Table S8.**
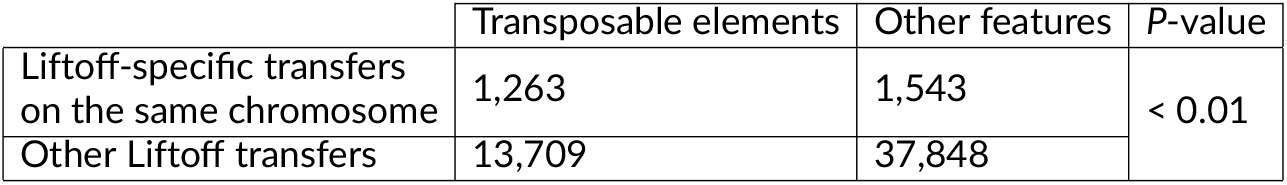
Transposable elements rates in Liftoff transfers. The *p*-value measures the enrichment in transposable elements in the Liftoff-specific transfers on the same chromosome, and was computed with Pearson’s Chi-squared test.

**Table S9.** Transposable elements rate in Liftoff transfers. The *p*-value measures the enrichment in transposable elements in the Liftoff-specific transfers in interchromosomal translocations, and was computed with Pearson’s Chi-squared test.

|  | Transposable elements | Other features | P-value |
| --- | --- | --- | --- |
| Liftoff-specific transfers on a different chromosome | 2,656 | 948 | < 0.01 |
| Other Liftoff transfers | 12,316 | 38,443 |  |

**Table S10.**
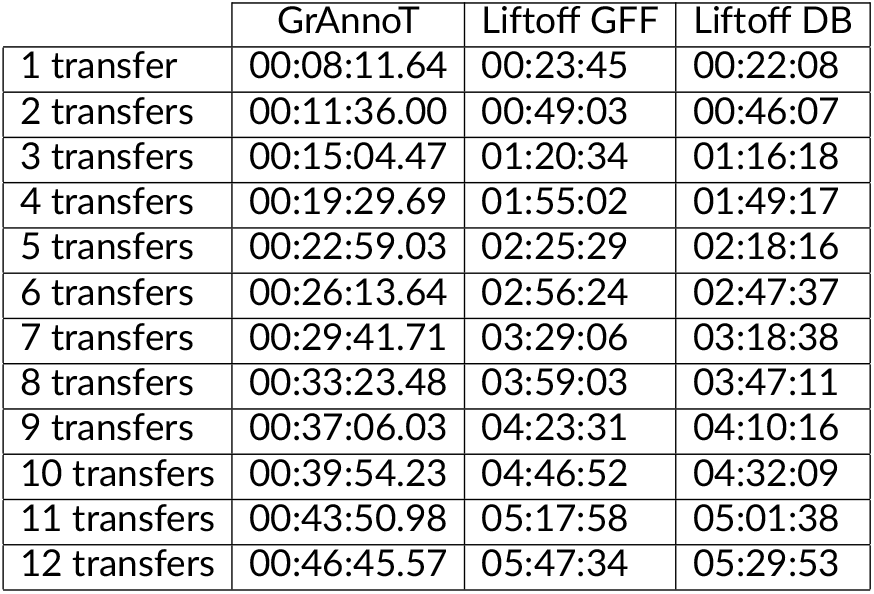
GrAnnoT and Liftoff CPU time comparison for 1 to 12 transfers.

**Table S11.** Indel rates in genes and CDS. Annotations were transferred between cv Nipponbare and cv Azucena, and the number of insertions and deletions was analyzed. The *p*-value measures the enrichment in indel in gene features, and was computed with Pearson’s Chi-squared test.

|  | Positions without indel | Positions with indel | P-value |
| --- | --- | --- | --- |
| Gene | 141,980,198 | 46,148 | < 0.01 |
| CDS | 74,201,787 | 8,762 |  |

**Table S12.** Frameshift indel rates in genes and CDS. Annotations were transferred between cv Nipponbare and cv Azucena, and the insertions and deletions lengths were analyzed. The *p*-value measures the enrichment in indel causing a frameshift in gene features, and was computed with Pearson’s Chi-squared test.

|  | Non-frameshift indel | Frameshift indel | P-value |
| --- | --- | --- | --- |
| Gene | 9,959 | 36,189 | < 0.01 |
| CDS | 3,950 | 4,812 |  |

**Table S13.**
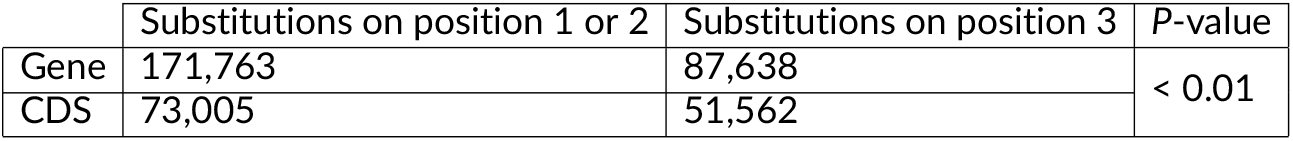
Substitution positions in nucleotide triplets in genes and CDS. Annotations were transferred between cv Nipponbare and cv Azucena, and the substitution positions were analyzed. The *p*-value measures the enrichment in substitutions on position 3 of the nucleotide triplets in CDS features, and was computed with Pearson’s Chi-squared test.

**Table S14.**
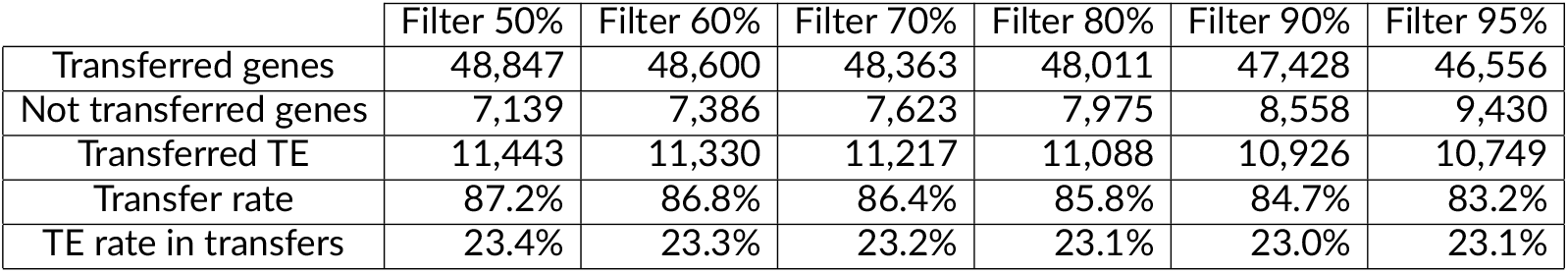
Impact of sequence identity and coverage based filtering on annotation transfers between cv Nipponbare and cv Azucena using GrAnnoT. The first row gives the value of both parameters. The filtering has a limited effect on gene transfer (transfer rate), and does not impact the transfer of transposable elements (TE).

